# Neural basis of opioid-induced respiratory depression and its rescue

**DOI:** 10.1101/2020.10.28.359893

**Authors:** Shijia Liu, Dongil Kim, Tae Gyu Oh, Gerald Pao, Jonghyun Kim, Richard D. Palmiter, Matthew R. Banghart, Kuo-Fen Lee, Ronald M. Evans, Sung Han

**Author notes:** Corresponding author: Sung Han, PhD. **Author Contributions:** S.H. conceived the idea, designed and constructed respiratory monitoring apparatus, and secured funding. S.H. and S.L. designed the experiments and wrote the manuscript with inputs from other authors. S.L. performed most of the experiments. D.K. and J.K. prepared the sample for RiboTag RNA sequencing. T.G.O. analyzed the data from RiboTag RNA sequencing. G.M.P. performed the CCM analysis. R.D.P. provided the Oprm1^*Cre*^ mouse line. M.R.B. provided the hMOR virus. K.F.L. and R.M.E. provided experimental resources.

## Abstract

Opioid-induced respiratory depression (OIRD) causes death following an opioid overdose, yet the neurobiological mechanisms of this process are not well understood. Here, we show that neurons within the lateral parabrachial nucleus that express the μ-opioid receptor (PBL^*Oprm1*^ neurons) are involved in OIRD pathogenesis. PBL^*Oprm1*^ neuronal activity is tightly correlated with respiratory rate, and this correlation is abolished following morphine injection. Chemogenetic inactivation of PBL^*Oprm1*^ neurons mimics OIRD in mice, whereas their chemogenetic activation following morphine injection rescues respiratory rhythms to baseline levels. We identified several excitatory G-protein coupled receptors expressed by PBL^*Oprm1*^ neurons and show that agonists for these receptors restore breathing rates in mice experiencing OIRD. Thus, PBL^*Oprm1*^ neurons are critical for OIRD pathogenesis, providing a promising therapeutic target for treating OIRD in patients.

## Introduction

The misuse of prescription and recreational opioids has taken nearly 4.5 million lives between 1999 and 2018 in the United States alone (1). The direct cause of death from opioid overdose is opioid-induced respiratory depression (OIRD) (2, 3). Currently, the only available antidote capable of reversing OIRD is naloxone, a nonselective opioid receptor antagonist. However, naloxone is associated with several disadvantages, namely the reappearance of OIRD due to its short half-life, the inability to reverse high-affinity opioid drugs (e.g., carfentanil and buprenorphine) due to its low binding affinity, and the potential to induce a catecholamine surge at high doses, which can cause cardiopulmonary arrest (4–6). Alternative non-opioid interventions include respiratory stimulants that act on the brainstem respiratory network (e.g., ampakines) or oxygen-sensing cells within the carotid bodies (e.g., K^+^-channel blockers) (5). However, none of these alternatives are as effective as naloxone, and the safety of these interventions has not been well documented. Therefore, novel treatment strategies are needed, and more effective countermeasures can only be developed with a mechanistic understanding of OIRD pathogenesis.

Neural substrates that contribute to OIRD pathogenesis should satisfy two criteria. They should localize to the breathing control network and express the μ-opioid receptor (MOR, encoded by the *Oprm1* gene), which has been demonstrated as the primary mediator of both the analgesic and respiratory effects of opioids (7, 8). Two candidate structures satisfy these criteria, namely the pre-Bötzinger complex (preBötC) and the parabrachial complex (9–17). The former lies in the ventrolateral medulla and is the primary generator of the respiratory rhythm (18–20). The parabrachial complex, including the lateral parabrachial, medial parabrachial, and Kölliker-Fuse (KF) nuclei, is located in the dorsolateral pons and modulates breathing (21–23), in response to homeostatic disturbances such as CO_2_/O_2_ imbalance and noxious stimuli (24–28). Recent studies have suggested the involvement of these two brain regions on OIRD pathogenesis by using either pharmacological or genetic approaches. Direct infusion of opioid agonists or antagonists into these brain areas induces or attenuates OIRD (9, 12, 29, 30). Region-specific genetic deletion of the *Oprm1* genes by viral delivery of Cre-recombinase into these areas of the *Oprm1*-floxed mice also attenuates OIRD (13, 14). However, the pharmacological approach lacks considerable specificity primarily due to the local diffusion of drugs and the difficulty of reproducible targeting, whereas the genetic deletion approach is often incomplete and prevents the possibility to directly control the neuronal activity in a spatially and temporally precise manner. As a result, discrepancies arise over the critical site of action responsible for OIRD (13, 14, 16, 17). To expand the scope of current research, our paper utilizes contemporary cell-type-specific approaches to characterize the role of a defined neuronal population in morphine-induced respiratory depression. This will allow us to better understand its cellular and physiological mechanisms and make inroads in developing therapeutic strategies.

Here we report that a genetically defined population of neurons that encodes the *Oprm1* gene in the lateral parabrachial nucleus (PBL^*Oprm1*^ neurons) is an important regulator of respiratory rhythm. Inhibition of these neurons by opioids leads to respiratory depression. Furthermore, we show that both chemogenetic activation of these neurons and pharmacological activation of endogenous G-protein coupled receptors (GPCRs) expressed specifically in these neurons completely rescues OIRD in mice.

## Results

To examine the contributions of global MOR signaling to OIRD, we generated an *Oprm1* reporter mouse line in which the endogenous *Oprm1* gene was knocked out by inserting a Cre:GFP cassette upstream of the start codon (*Oprm1^Cre^*) (Figure 1B and Supplementary Figure 1A for validation), and measured the effects of systemic morphine injection on breathing rhythms via whole-body plethysmography (Figure 1A). After receiving a morphine injection, wild-type mice exhibited a significantly lower respiratory rate (46.5 ± 3.1% of the pre-morphine baseline (data are represented as mean ± SEM throughout the paper)). In contrast, mice lacking one or both functional copies of the *Oprm1* gene (*Oprm1^Cre/+^* and *Oprm1^Cre/Cre^*) exhibited more moderate reductions in respiratory rate (63.2 ± 2.8% and 91.2 ± 2.8% of the pre-morphine baseline, respectively) (Figure 1C, D). *Oprm1*, therefore, plays a vital role in OIRD.

**Figure 1.**
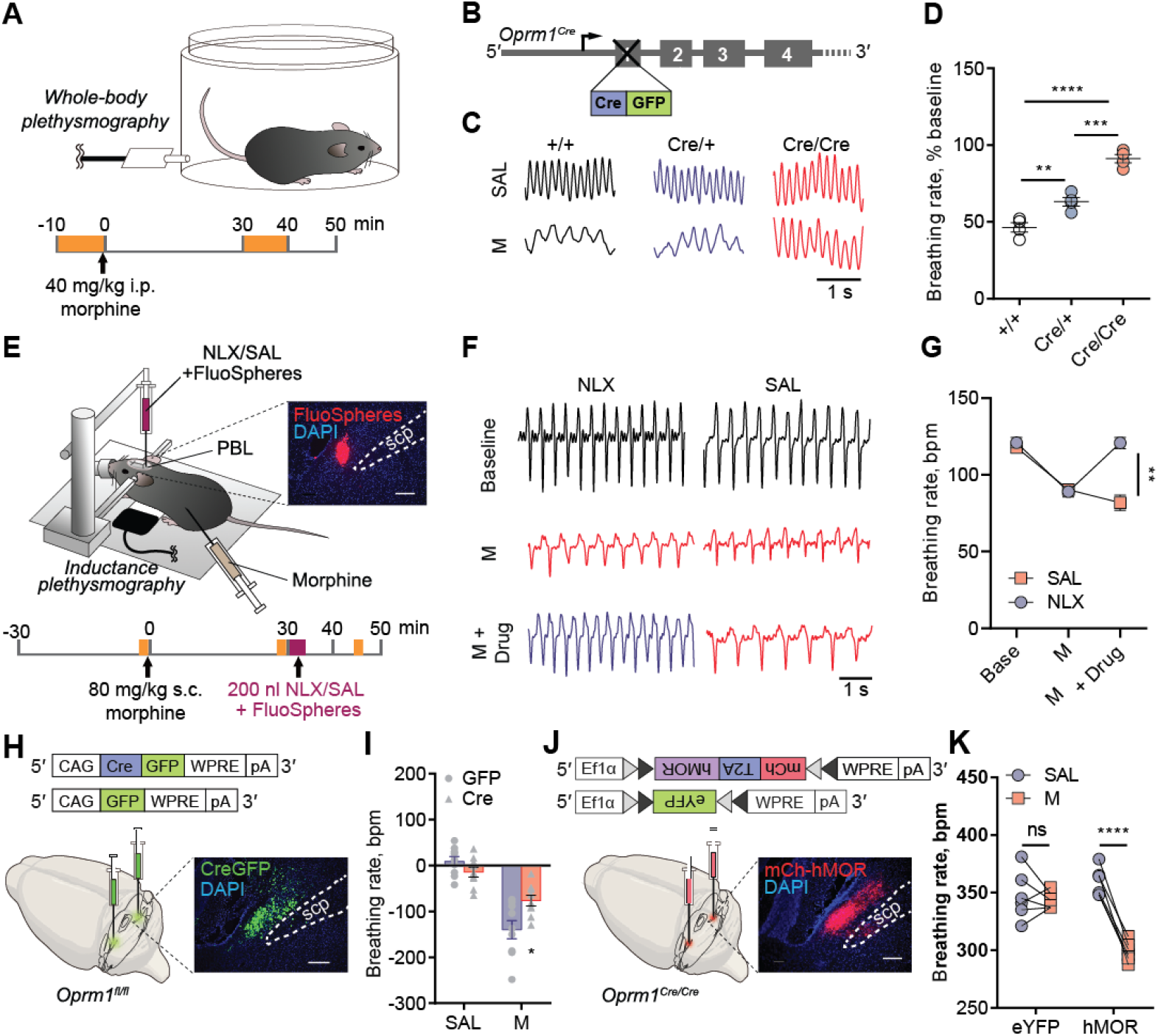
MOR signaling in the PBL mediates morphine-induced respiratory depression. (**A**) Schematics of breathing monitoring in awake mice with whole-body plethysmography before and after systemic morphine injection (refer to **C**, **D**, **I**, **K**). (**B-D**) *Oprm1^+/+^*, *Oprm1^Cre/+^*, and *Oprm1^Cre/Cre^* mice (**B**) displayed different breathing patterns (**C**) and rate changes (**D**) in response to systemic morphine injection. (**E**) Schematics of breathing monitoring in anesthetized mice with inductance plethysmography, during systemic morphine injection and PBL-specific blockade of MOR signaling. Onset, histology of the co-injected FluoSpheres in the PBL. Scp, superior cerebellar peduncle. (**F**) Representative plethysmographs from naloxone (NLX) and saline (SAL) injected groups before morphine, after morphine, and after drug injections into the PBL. (**G**) Injecting naloxone (n = 5), but not saline (n = 5) into the PBL rescued morphine-induced respiratory depression. (**H**) Schematics of PBL-specific knockout of the *Oprm1* gene by stereotaxically injecting AAV-Cre-GFP into the PBL of the *Oprm1^fl/fl^* mice. (**I**) PBL-specific *Oprm1* knockout (n = 9) significantly attenuated OIRD compared to the GFP injected control group (n = 9) after systemic morphine injection. (**J**) Schematics of PBL-specific rescue of the *Oprm1* gene in *Oprm1*-null background by stereotaxically injecting AAV-DIO-hMOR into the PBL of the *Oprm1^Cre/Cre^* mice. (**K**) Mice with PBL-specific rescue of *Oprm1* (n = 6) displayed OIRD phenotype, whereas eYFP injected control group (n = 6) remained insensitive after systemic morphine injection. Data are presented as mean ± SEM. Statistical tests, One-way ANOVA with Tukey’s multiple comparison post-hoc test (**D**) or Two-way ANOVA with Bonferroni’s multiple comparison post-hoc test (**G, I, K**). *, *p* < 0.05; **, *p* < 0.01; ***, *p* < 0.001; ****, *p* <0.0001. Scale bar, 1s (**C, F**) or 200 μm (**E, H, J**).

We then used pharmacological and genetic tools to manipulate opioid signaling specifically in the PBL. First, wild-type mice received a systemic injection of morphine followed by an infusion of naloxone directly into the PBL (Figure 1E). To facilitate stereotaxic naloxone delivery, we lightly anesthetized mice with 1-1.5% isoflurane and monitored their breathing using a piezoelectric sensor that detects chest movements (31). After systemic injection of morphine, the breathing rate dramatically decreased from 121.0 ± 3.4 bpm to 89.1 ± 3.5 bpm, and subsequent stereotaxic injection of naloxone into the PBL recovered the breathing rate back to pre-morphine baseline (120.8 ± 3.7 bpm) (Figure 1F, G). We then explored whether the expression of the *Oprm1* gene in the PBL is necessary and sufficient for OIRD. To assess necessity, we conditionally knocked out the *Oprm1* gene using stereotaxic delivery of a recombinant adeno-associated virus (AAV) expressing Cre recombinase into the PBL of the *Oprm1^fl/fl^* mice, in which the *Oprm1* gene is flanked by loxP sites (Figure 1H). Compared to mice receiving GFP-expressing control virus, PBL-specific knockout of the *Oprm1* gene attenuated OIRD following systemic injection of morphine, as measured by whole-body plethysmography (breathing rate changes in GFP group: −136.6 ± 21.1 bpm; Cre group: −76.4 ± 11.6 bpm) (Figure 1I and Supplementary Figure 2A-C). To test sufficiency, we reintroduced the wild-type *Oprm1* gene into the PBL^*Oprm1*^ neurons of Cre-expressing *Oprm1*-null mice. This was achieved by injecting an AAV expressing Cre-dependent human MOR (AAV-hSyn-DIO-mCherry-T2A-FLAG-hMOR) (Figure 1J) into the PBL of *Oprm1^Cre/Cre^* mice. Selective expression of hMOR in the PBL restored the OIRD phenotype (hMOR group, post-saline: 359.0 ± 5.1 bpm; post-morphine: 299.0 ± 3.6 bpm) compared to the control group (Figure 1K and Supplementary Figure 2D). Note that the breathing rate measured by the piezoelectric sensor under anesthesia (Figure 1G) is ~3-fold lower than that measured in a plethysmography chamber (Figure 1K). Together, these data strongly support the conclusion that the *Oprm1* gene expression in the PBL mediates OIRD in mice.

We then focused on PBL^*Oprm1*^ neurons and explored their relationship with OIRD. PBL^*Oprm1*^ neurons are mostly glutamatergic and express higher levels of the *Oprm1* gene than the adjacent KF nucleus, which provides post-inspiratory drive (13, 23, 31, 32) (Supplementary Figure 1B-C). We devised an experimental platform to simultaneously measure PBL^*Oprm1*^ neuronal activity and respiration in awake, freely moving mice using fiber-photometry calcium imaging *in vivo* with a thermistor implanted into the nasal cavity (Figure 2A). The thermistor-based respiration monitoring device detects the temperature change between inhaled and exhaled air and converts breath-to-breath fluctuations in resistance into voltage signals (33). We stereotaxically injected an AAV expressing Cre-dependent jGCaMP7s (AAV-DIO-jGCaMP7s) into the PBL, then implanted an optic fiber into the PBL of *Oprm1^Cre/+^* mice and a respiration sensor in the nasal cavity (Figure 2B). PBL^*Oprm1*^ activity and breathing rate exhibited a tight positive correlation (Figure 2C-D, H). Moreover, morphine administration substantially decreased the breathing rate (46.5 ± 3.9% of the pre-morphine baseline) (Figure 2H, J, L, and N) despite the potent effect morphine has on locomotor sensitization (Supplementary Figure 3), which one would expect to elevate the breathing rate. Besides, morphine decreased calcium activity (Figure 2H, J, O, and Q), as well as eliminated any spontaneous fluctuations in both breathing and calcium signals (Figure 2N, Q). In contrast, saline injection largely preserved the original patterns of the signals (Figure 2D, 2F, 2L-M, and 2O-P). Using convergent cross-mapping (CCM) (34), a statistical algorithm that makes predictions from independent time-series data, we predicted changes in respiratory rate using the calcium signal as input (76.2 ± 8.1%, pre-morphine predictability) (Supplementary Figure 4). Interestingly, systemic morphine administration not only reduced the correlation coefficient (Figure 2I, K) but also completely abolished the predictive relationship between respiratory rate and the calcium signal (12.5 ± 20.2%, post-morphine predictability) (Supplementary Figure 4C-D). Conversely, the saline injection did not alter the correlation between these signals (Fig 2E, G, and Supplementary Figure 4A-B). These results indicate that PBL^*Oprm1*^ neurons are critically involved in modulating respiratory rate and that morphine suppresses both the PBL^*Oprm1*^ neuronal activity and its coupling with the breathing rate.

**Figure 2.**
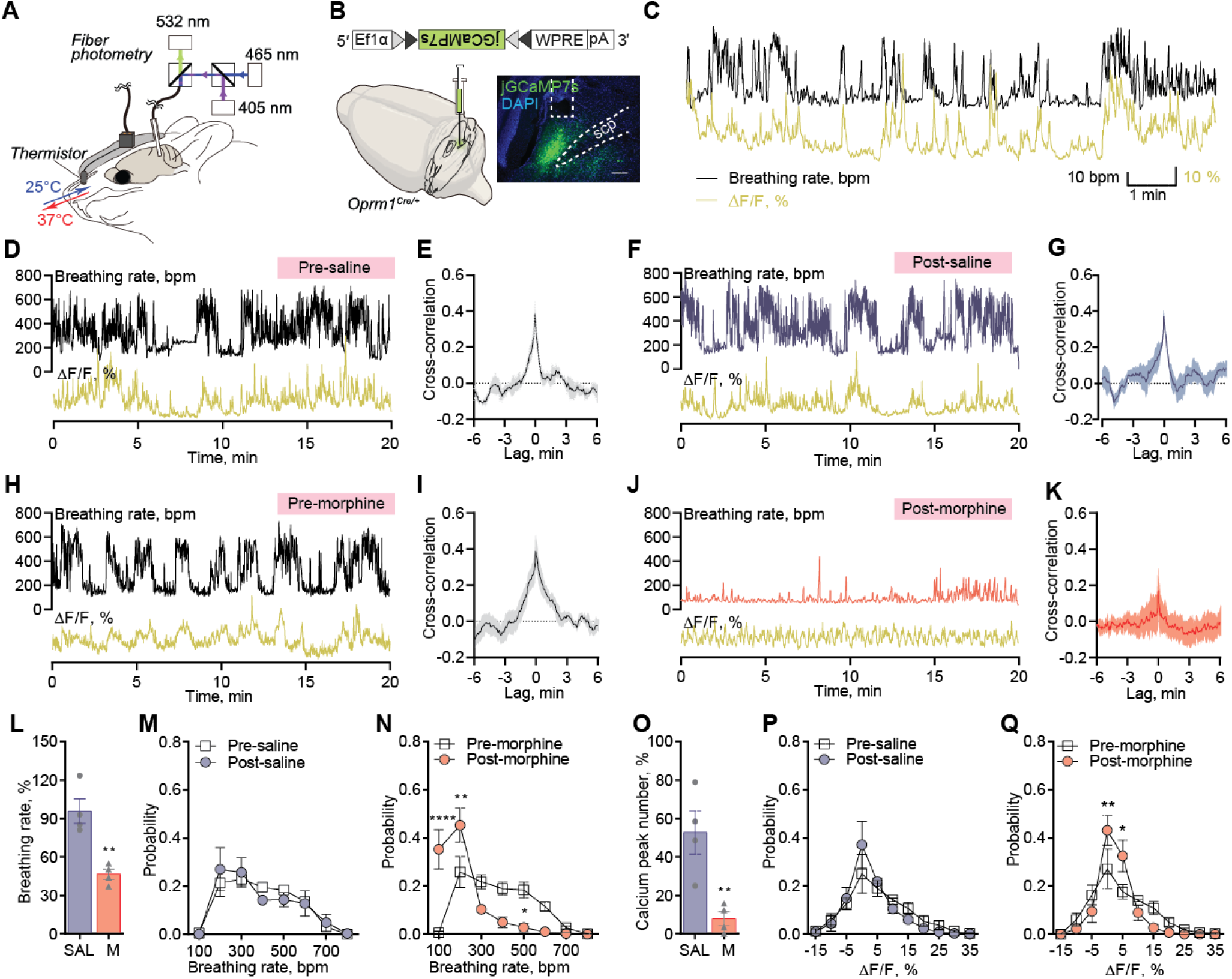
Morphine disrupts the tight correlation between PBL ^*Oprm1*^ neuronal activity and respiratory rate. (**A**) Schematics of simultaneous recording of respiration and PBL^Oprm1^ calcium activity with a thermistor sensor (implanted in the nasal cavity) and fiber photometry. (**B**) The genetically encoded calcium indicator, jGCaMP7s, was specifically expressed in the PBL^*Oprm1*^ neurons by stereotaxic injection of AAV-DIO-jGCaMP7s into the PBL of the *Oprm1^Cre/+^* mice. (**C**) Time-matched traces of PBL^*Oprm1*^ activity and breathing rate as calculated from the thermistor sensor. (**D, F, H, J**) Simultaneously recorded PBL^*Oprm1*^ calcium activity and respiratory rate before and after systemic saline or morphine injection. (**E, G, I, K**) Cross-correlogram of calcium signal and respiratory rate in the pre-saline, post-saline, pre-morphine, and post-morphine groups (n = 4). (**L**) Morphine injection significantly depressed respiratory rate to around 50% of the baseline (n = 4), whereas saline injection (n = 4) did not affect the respiratory rate. (**M-N**) Normalized distribution of breathing rate before and after saline (**M**) and morphine (**N**) injections (n = 4). (**O**) The number of calcium peaks was significantly decreased after morphine injection (n = 4) compared to saline injection (n = 4). (**P-Q**) Normalized distribution of calcium signals before and after saline (**P**) and morphine (**Q**) injections (n = 4). Data are presented as mean ± SEM. Statistical tests, Unpaired t-test (**L, O**) or Two-way ANOVA with Bonferroni’s multiple comparison post-hoc test (**M, N, P, Q**). *, *p* < 0.05; **, *p* < 0.01; ****, *p* < 0.0001. Scale bar, 200 μm.

To further investigate PBL^*Oprm1*^ neurons’ role in breathing regulation, we specifically manipulated their activity using chemogenetic tools, namely Designer Receptor Exclusively Activated by Designer Drugs (DREADD) (35). If PBL^*Oprm1*^ neurons directly regulate breathing, inhibiting their activity via a G_i/o_-coupled DREADD should recapitulate the OIRD phenotype. Furthermore, this should also happen in *Oprm1* knockout mice, which failed to display OIRD when given systemic morphine (Figure 1C-D). We expressed a κ-opioid receptor-derived DREADD (KORD) (36) Cre-dependently and bilaterally in PBL^*Oprm1*^ neurons of *Oprm1^Cre/Cre^* mice (Figure 3A). Systemic injection of its synthetic ligand, salvinorin B (SALB), decreased the respiratory rate (317.8 ± 3.6 bpm) compared to the DMSO-injected control group (342.4 ± 4.1 bpm) (Figure 3B and Supplementary Figure 5A-B). We did not observe significant breathing amplitude changes upon SALB injection in either control or KORD-expressing animals (Supplementary Figure 5B-C).

**Figure 3.**
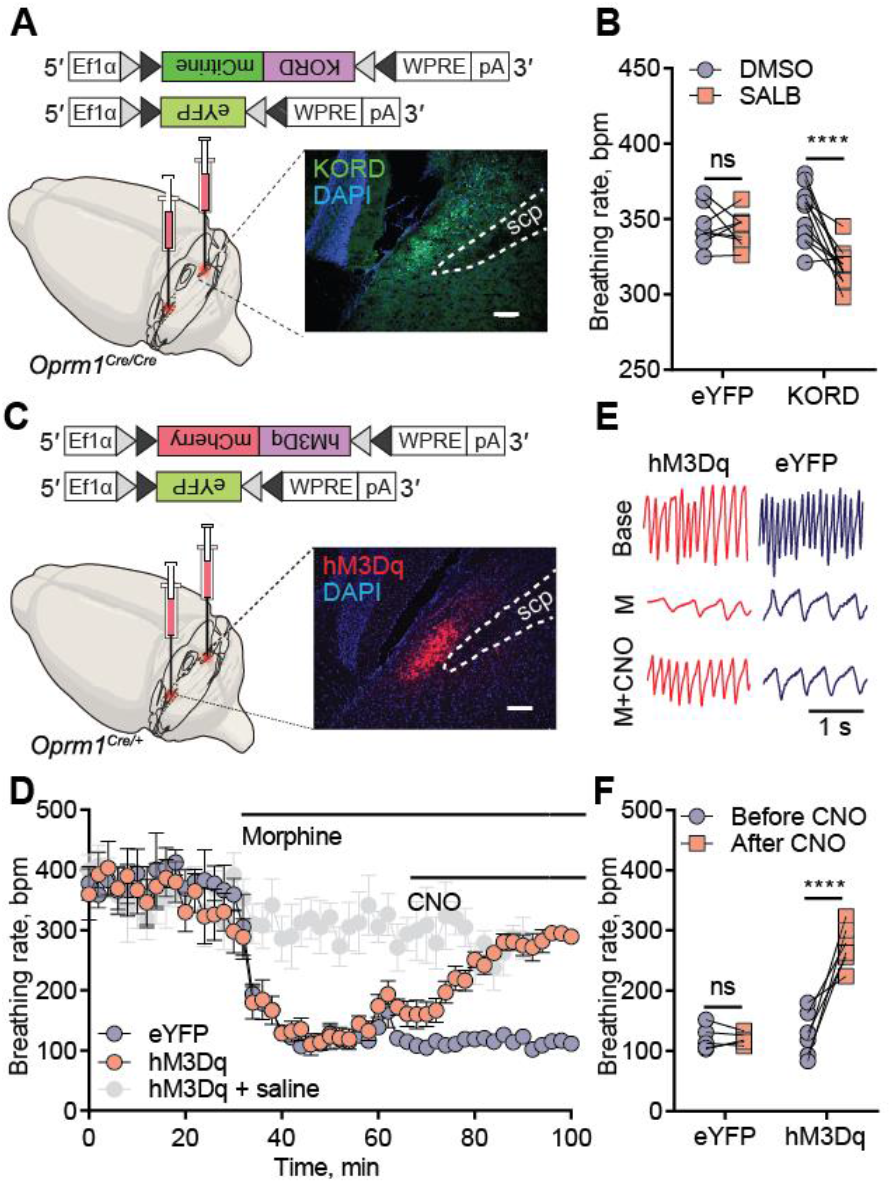
Inhibition of PBL^*Oprm1*^ neurons mimics OIRD, and chemogenetic activation of PBL^*Oprm1*^ neurons rescues OIRD. (**A**) Schematics of mimicking OIRD through PBL^*Oprm1*^ inhibition using an inhibitory κ-opioid receptor-derived DREADD (KORD). Here, AAV-DIO-KORD-mCitrine is stereotaxically injected into the bilateral PBL of *Oprm1^Cre/Cre^* mice to express in the PBL^*Oprm1*^ neurons. (**B**) Salvinorin B (SALB, 7.5 mg/kg) injection significantly decreased respiratory rate compared to the DMSO control in the KORD-expressing mice (n = 11). In contrast, no changes were observed in eYFP-expressing mice (n = 8). (**C**) Schematics of rescuing OIRD through PBL^*Oprm1*^ activation by an excitatory DREADD, hM3Dq. Here, AAV-DIO-hM3Dq-mCherry was expressed in the bilateral PBL of *Oprm1^Cre/+^* mice. (**D**) Activation of PBL^*Oprm1*^ neurons by injecting Clozapine N-oxide (CNO, 5 mg/kg) rescued the morphine-induced respiratory depression to control levels in the hM3Dq-expressing group (n = 6), but not the eYFP-expressing group (n = 5). (**E-F**) Representative plethysmograph and quantitative analysis depicting OIRD rescued by CNO injection in the hM3Dq-expressing group (n = 7), but not in the eYFP-expressing group (n = 5). Statistical tests, Two-way ANOVA with Bonferroni’s multiple comparison post-hoc test. ns, not significant; ****, *p* < 0.0001. Scale bar, 200 μm.

We next asked whether artificial activation of PBL^*Oprm1*^ neurons could prevent OIRD in mice. We expressed hM3Dq, a metabotropic acetylcholine receptor-derived excitatory DREADD, in PBL^*Oprm1*^ neurons via bilateral injection of an AAV encoding Cre-dependent hM3Dq into *Oprm1^Cre/+^* mice (Figure 3C). Additionally, we implanted a micro-thermistor sensor into the nasal cavity to monitor respiratory rhythm in awake, freely moving mice over the course of OIRD (Supplementary Figure 5D). Upon systemic morphine injection, the respiratory rate decreased significantly within 10 min (from 332.0 ± 25.6 bpm to 124.6 ± 9.1 bpm) (Figure 3D-E). We then systemically injected the hM3Dq agonist, clozapine-N-oxide (CNO), into the same animal to activate PBL^*Oprm1*^ neurons. CNO injection increased the breathing rate in hM3Dq-expressing animals (268.0 ± 13.5 bpm) to a level comparable to controls that did not receive morphine (275.7 ± 24.9 bpm). Nevertheless, CNO did not rescue breathing rates in eYFP-expressing control animals (120.0 ± 4.3 bpm) (Figure 3D-F and Supplementary Figure 5E). A complete rescue was also observed in minute volume (Supplementary Figure 5F, G). These data show that activation of PBL^*Oprm1*^ neurons is sufficient to restore normal breathing rhythms in mice displaying OIRD.

Having determined that chemogenetic activation of PBL^*Oprm1*^ neurons could effectively prevent OIRD, we sought to identify endogenous signaling pathways that could be used to activate PBL^*Oprm1*^ neurons. To identify the active transcriptome enriched in PBL^*Oprm1*^ neurons, we sequenced ribosome-associated mRNAs from the PBL of *Oprm1^Cre^*::RiboTag mice, which express hemagglutinin (HA)-tagged ribosomal protein Rpl22 in *Oprm1^+^* neurons (Figure 4A-B) (37). We identified 69 mRNAs upregulated over three folds in PBL^Oprm1^ neurons than non-*Oprm1* neurons in the PBL (Figure 4C). *Oprm1*, as well as other major parabrachial markers (*Tac1, Foxp2*, and *Adcyap1*) were enriched, but *Calca* and *Pdyn*, markers for the external lateral and dorsal lateral parabrachial nucleus, were not (38, 39). Notably, the PBL^*Oprm1*^ active transcriptome revealed several GPCRs expressed at high levels in these neurons, which we subsequently investigated as potential pharmacological targets to rescue OIRD.

**Figure 4.**
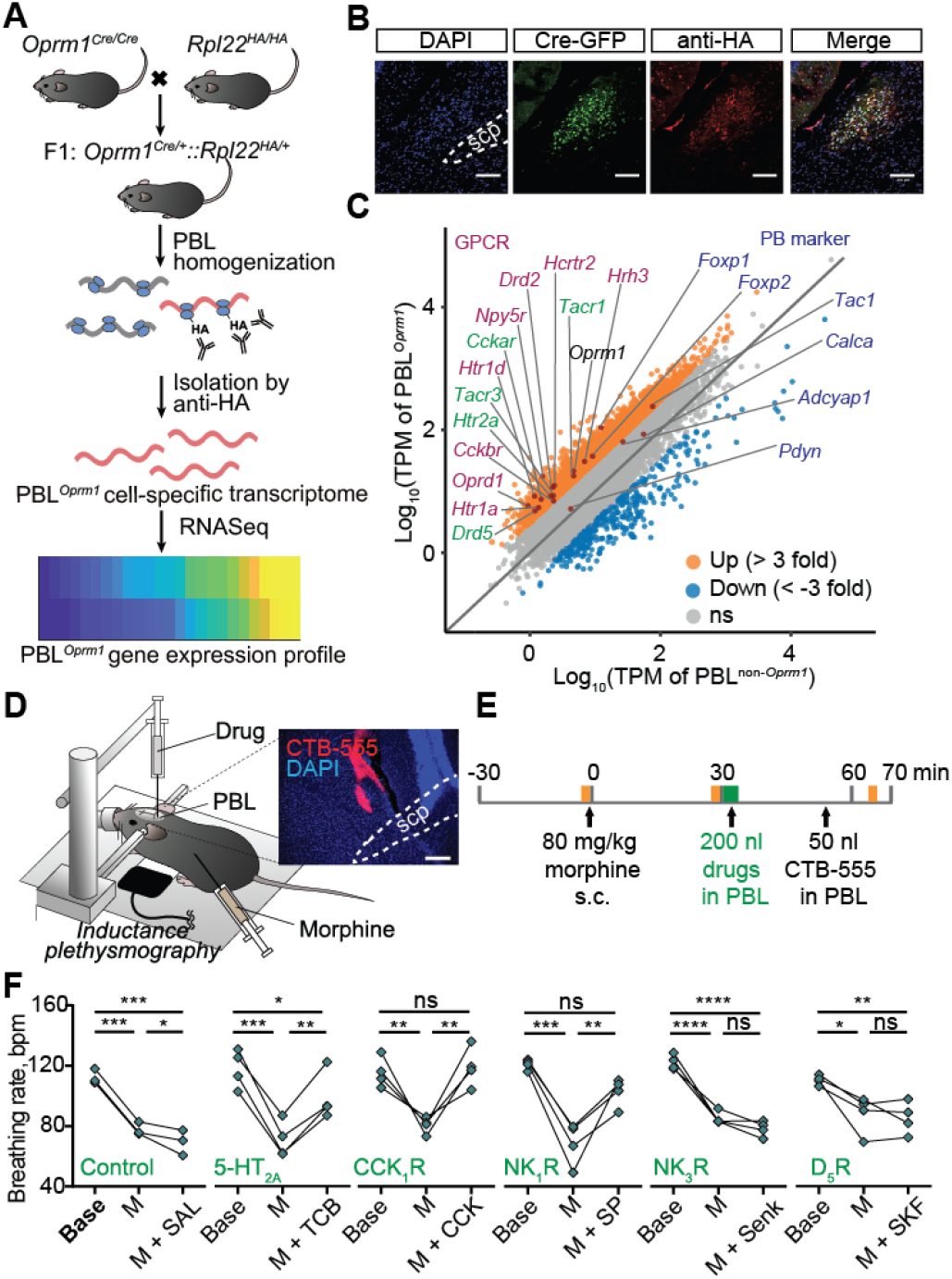
Rescue of OIRD through PBL^*Oprm1*^ stimulation via receptors identified by RiboTag RNA sequencing. (**A**) Schematics of obtaining PBL^*Oprm1*^ active transcriptome. *Oprm1^Cre/Cre^* mice were crossed with RiboTag mice, which express hemagglutinin-tagged ribosomal protein L22 (*Rpl22^HA^*) in a Cre-dependent manner. After collecting the PBL, active mRNAs bound to the *Rpl22^HA^* were isolated with anti-HA antibody-mediated ribosome pull-down and sequenced. (**B**) Immunohistochemistry of the PBL region of the *Oprm1^Cre^*::*Rpl22^HA^* mice with anti-GFP and anti-HA antibodies showed that the Cre:GFP and Rpl22^HA^ were co-expressed in PBL^*Oprm1*^ neurons. (**C**) Fourteen GPCR mRNAs (magenta and green) were enriched more than three-fold in PBL^*Oprm1*^ neurons compared with non-*Oprm1* neurons in the PBL. Conventional parabrachial markers (dark blue), such as *Foxp2*, *Tac1*, and *Adcyap1* genes were enriched in PBL^*Oprm1*^ neurons, but not *Pdyn* and *Calca*. (**D-E**) Schematics of rescuing OIRD through artificial stimulation of receptors expressed on PBL^*Oprm1*^ neurons by PBL-specific injection of the agonists. Onset, histology of cholera toxin subunit B (CTB)-555 for marking the injection site. (**F**) Breathing rate decreased upon systemic morphine injection and rescued by PBL-specific injection of the agonists of 5-HT_2A_, CCK_1_R, and NK_1_R. For the list of pharmacological agents, see Supplementary Table 1. Statistical tests, RM One-way ANOVA with Tukey’s multiple comparison post-hoc test, ns, not significant; *, *p* < 0.05; **, *p* < 0.01; ***, *p* < 0.001, ****, *p* < 0.0001. Scale bar, 100 μm (**B**) or 200 μm (**D**).

We selected five excitatory GPCRs for pharmacological manipulation and confirmed that they were expressed in PBL^*Oprm1*^ neurons by RNA *in situ* hybridization (Supplementary Figure 6A-B). These GPCRs were 5-hydroxytryptamine receptor 2A (*Htr2a*), cholecystokinin A receptor (*Cckar*), tachykinin receptor 1 (*Tacr1*), tachykinin receptor 3 (*Tacr3*), and dopamine receptor D5 (*Drd5*). We systemically injected their agonists (TCB-2, CCK8S, Substance P, Senktide, and SKF83959, respectively) into the PBL of anesthetized mice after inducing OIRD (breathing rate before morphine: 115.4 ± 1.6 bpm, after morphine: 77.9 ± 2.3 bpm; Figure 4D-E and Supplementary Table 1). Three of the five agonists tested (TCB-2, CCK8S, and Substance P) increased the respiratory rate (98.1 ± 7.9, 117.1 ± 6.4, 100.8 ± 4.6 bpm; Figure 4F and Supplementary Figure 6C-E). Notably, these same drugs have been shown to modulate respiratory rhythm in different contexts by activating other respiratory centers, and some have been associated with an anti-morphine effect (40–42). Our findings suggest that activating PBL^*Oprm1*^ neurons through endogenous signaling pathways may effectively treat OIRD in patients.

## Discussion

We have identified a population of neurons in the pontine respiratory group, the PBL^*Oprm1*^ neurons, as critical mediators of morphine-induced respiratory depression and candidates for its rescue in intact mice. Genetic deletion of *Oprm1* from the PBL attenuated OIRD, while reintroducing human MOR into the PBL^*Oprm1*^ neurons restored OIRD. By combining cell-type-specific tools with breathing monitoring in awake behaving mice, we have demonstrated that PBL^*Oprm1*^ neurons are regulators of the breathing rhythm and pattern. Systemic morphine administration dramatically abolishes the activity of these neurons and weakens their control on breathing. Furthermore, chemogenetic inhibition of PBL^*Oprm1*^ neurons mimics OIRD, and prolonged optogenetic inhibition of PBL^*Oprm1*^ neurons induces apnea (observations from our group (43)), which precedes cardiorespiratory collapse prior to overdose-induced death. By contrast, artificial activation of PBL^*Oprm1*^ neurons, either chemogenetically or pharmacologically, successfully restores breathing in mice experiencing OIRD (Supplementary Figure 7).

### Investigating OIRD with cell-type-specific approaches

Recent studies have demonstrated the involvement of both pontine and medullary respiratory groups in OIRD (13, 14) without reaching a consensus on the key players. We think this is mostly due to the lack of specificity from the pharmacological and genetic deletion approaches and the intricacies of physiological preparations, both of which hampers reproducibility. In comparison, we exploited the *Oprm1^Cre^* mouse line which provides access to a restricted neuronal population, and obtained multi-faceted evidence supporting the role of the parabrachial nucleus in OIRD. In our view, OIRD is a synergistic outcome from impairment of the interconnected pontomedullary breathing network that widely expresses MOR both pre- and post-synaptically (10, 11, 30, 44, 45). For example, parabrachial neurons send tonic excitatory inputs to the preBötC (39, 43, 46, 47), and both of these areas express MOR (18, 48).

One challenge of elucidating the OIRD mechanism is how to access a group of neurons while simultaneously record breathing over the course of OIRD. Traditional respiratory monitoring approaches in rodents, such as *in situ* or *in vivo* anesthetized preparations and plethysmography chambers, are more challenging to achieve this goal. Therefore, we utilized the implantable temperature sensor to monitor breathing in awake, unrestrained, behaving mice with unprecedented freedom to access the brain due to its open configuration. With this approach, we implemented chemogenetics and *in vivo* calcium imaging while simultaneously monitored breathing rhythms. Similar techniques can be applied to the *Oprm1*-expressing neurons in the preBötC, KF, the postinspiratory complex (49), and other respiratory centers to resolve the long-standing debate on their contributions in OIRD (16, 17).

### PBL^*Oprm1*^ neurons’ role in breathing regulation and OIRD

The parabrachial complex has long been reported to modulate respiration and sometimes exert opposing effects depending on the location of artificial manipulation (21–23). Stimulation of the PBL results in tachypnea, whereas stimulation of the KF elicits prolonged post-inspiration and bradypnea (13, 31, 32). Consistent with these previous reports, our monitoring and manipulation data suggest that PBL^*Oprm1*^ neurons have the most substantial effect on the breathing rate (23). In addition to breathing, the PBL also regulates various homeostatic functions (39) and incorporates the alarm center, which senses deviations from the homeostasis and relays aversive signals to higher-order brain structures (38, 50). At the same time, by sending tonic excitatory outputs to medulla respiratory centers (39, 43, 46), the parabrachial neurons can increase breathing rhythm under conditions that immediately require more oxygen due to metabolic needs, such as hypoxia (27) and hypercapnia (26), in particular, hypercapnic arousal during sleep (24, 28, 51); or non-metabolic behavioral demands, such as escaping from a threat (25). As a representative glutamatergic population in the PBL (Supplementary Figure 1), *Oprm1*-positive neurons are likely to participate in similar physiological processes. Our data demonstrated that an overdose of morphine shuts down PBL^*Oprm1*^ neurons’ activity, preventing them from responding to conditions that decrease respiratory rhythm. These conditions may include hypoxia, hypercapnia, anesthesia, or sedative drug treatment. Indeed, the incidence of OIRD is dramatically increased when opioids are used together with sedatives and anesthetics (52), and the failure to increase respiratory behavior in response to hypercapnic gas is one characteristic of OIRD (3). On the contrary, a mild inhibition of PBL^*Oprm1*^ neurons may explain many of the behavioral effects of lower doses of endogenous or exogenous opioids, such as slowed breathing, calmness, and reward (43). The current study used morphine as a starting point to induce OIRD, and further investigation is required to determine the involvement of PBL^*Oprm1*^ neurons under the challenge of other types of sedative agents.

### A proof of principle of rescuing OIRD by activating a pontine population

PBL^*Oprm1*^ neurons are among the key nodes most vulnerable to opioid action, and their activation is sufficient to restore the breathing rate and tidal volume following morphine administration. This activation can be achieved by exploiting both artificial and, more importantly, endogenous signaling pathways. Besides, manipulating PBL^*Oprm1*^ neurons located in the dorsolateral pons is associated with lower risk than the respiratory rhythm generators within the deep medulla. We have provided a proof of concept of rescuing OIRD by activating endogenous G_q/s_-coupled GPCRs expressed in PBL^*Oprm1*^ neurons (Supplementary Figure 7). In principle, cell-type-specific manipulation can also be achieved through the collective activation of multiple GPCRs.

Our transcriptomic profiling data suggested that other neuromodulators, such as serotonin, cholecystokinin, and tachykinin, may also involve in respiratory regulation with potential interplay with the opioid system (40–42). The same receptor that we identified, Htr2a, has recently been confirmed to express in the PBL and mediate hypercapnia-induced arousal by integrating serotonergic inputs from the dorsal raphe (51). It would be interesting to identify other neuromodulatory circuits that may be involved in the regulation of OIRD. Similar transcriptomic profiling strategies can also be applied to the broader breathing control network to exploit the therapeutic potential of GPCRs in reversing OIRD.

## Acknowledgments

We thank the Han lab members for the critical discussion of the paper and D. O’Keefe and M. Shields for critical inputs on the manuscript. S.H. is supported by 1R01MH116203 from NIMH, the Bridge to Independence award from the Simons Foundation Autism Research Initiative (SFARI #388708), and Brain Research Foundation Fay/Frank Seed Grant. K.F.L is supported by MH114831, OD023076, AG062232, NS115183 and AG064049.

## Supplementary Materials

### Materials and Methods

#### Experimental Animals

The *Oprm1^Cre:GFP^* mouse line was generated from the lab of Dr. Richard Palmiter (see below). C57BL/6J (Stock No. 000664), *Oprm1^fl/fl^* (Stock No. 030074), RiboTag *Rpl22^HA/HA^* (Stock No. 011029) and Ai14 *Gt(ROSA)26Sor^tm14(CAG-tdTomato)Hze^* (Stock No. 007914) mouse lines were obtained from The Jackson Laboratory. Homozygous RiboTag and Ai14 mice were crossed with homozygous *Oprm1*^Cre^ mice for RiboTag and anatomy experiments, respectively. Both male and female mice were used in all studies. Animals were randomized to experimental groups, and no sex differences were noted. Mice were maintained on a 12-h light/dark cycle and provided with food and water *ad libitum*.

#### Generation of *Oprm1^Cre^* mice

A cassette encoding Cre:GFP was inserted just 5’ of the initiation codon in the first coding exon of the *Oprm1* gene. The 5′ arm (~3.1 kb with *Pac*I and *Sal*I sites at 5’ and 3’ ends, respectively) and 3′ arm (~4.5 kb with *Xho*I and *Not*I sites at 5’ and 3’ ends, respectively) of *Oprm1* gene were amplified from a C57BL/6 BAC clone by PCR using Q5 Polymerase (New England Biolabs, USA) and cloned into polylinkers of a targeting construct that contained mnCre:GFP, a FRT-flanked Sv40Neo gene for positive selection, and HSV thymidine kinase and *Pgk*-diphtheria toxin A chain genes for negative selection. The mnCre:GFP cassette has a Myc-tag and nuclear localization signals at the N-terminus of Cre recombinase, which is fused to green fluorescent protein followed by a SV40 polyadenylation sequence. The construct was electroporated into G4 ES cells (C57BL/6 × 129 Sv hybrid), and correct targeting was determined by Southern blot of DNA digested with *Kpn*I using a ^32^P-labeled probe downstream of the 3′ arm of the targeting construct. Twelve of the 84 clones analyzed were correctly targeted. One clone that was injected into blastocysts resulted in good chimeras that transmitted the targeted allele through the germline. Progeny were bred with *Gt(Rosa)26Sor-FLP* recombinase mice to remove the SV-Neo gene. Mice were then continuously backcrossed to C57BL/6 mice. Routine genotyping is performed with three primers: 5’ CCT TCC ACT CAG AGA GTG GCG (*Oprm1* forward), 5’ CCT TCC ACT CAG AGA GTG GCG (*Oprm1* reverse), and 5’ GGC AAA TTT TGG TGT ACG GTC AG (Cre reverse). The wild-type allele gives a band of ~500 bp, while the targeted allele provides a band with ~400 bp after 34 cycles with 20-s annealing at 60 °C.

#### Respiratory measurements

##### Inductance plethysmography

Inductance plethysmography was performed by placing a piezoelectric film beneath the chest of an anesthetized animal, which converts the chest movement into a voltage signal. The PowerLab system with LabChart 8 software (ADInstruments Inc., USA) was used for data acquisition, inspiratory and expiratory peak detection, and rate and amplitude calculation. Data were sampled at 100 or 400 Hz, low-pass filtered at 10 Hz, and smoothed with a 100-ms moving window. Automatic peak detection was validated with manual peak detection. Since the location of the sensor is subject to the slight movements of the animal’s body, the raw voltage value of the respiratory peak is less representative of the breathing amplitude.

##### Whole-body plethysmography (WBP)

A custom-built WBP chamber was utilized for measuring respiratory changes. The PowerLab system with LabChart 8 software was used for data acquisition, inspiratory and expiratory peak detection, and rate and amplitude calculation. Data were sampled at 1 kHz, band-pass filtered at 1–10 Hz, and smoothed with a 100-ms moving window. Automatic peak detection was validated with manual peak detection.

Mice were introduced into the WBP chamber for three 20 min habituation sessions before testing. Mice were kept in the chamber for 10 - 12 min during the testing session before and after the drug injection. After 5 - 10 min of chamber introduction, a stable pattern was reached, and the averaged value of a stabilized 1 min segment was analyzed.

##### Micro thermistor-based plethysmography

A custom-built micro thermistor was implanted into the mouse nasal cavity to detect changes in temperature between inspiratory and expiratory airflow (*18*). The sensor was assembled using a Negative Temperature Coefficient (NTC) thermistor (TE Connectivity Ltd., Switzerland), an interconnector (Mill-Max Mfg. Corp., USA), and a voltage divider (Phidgets Inc., Canada). PowerLab was used for data acquisition, inspiratory and expiratory peak detection, and rate and amplitude calculation. Data were sampled at 1 kHz, filtered with a 0.4 - 25 Hz band-pass filter, and smoothed with a 50-ms moving window. Automatic peak detection was validated with manual peak detection.

Minute volume is approximated by first calculating the integral of the voltage channel for each breath, summing across one minute, then normalizing against the average value of the first 30 minutes. Values are shown after normalization as the raw values vary considerably across animals. For comparison before and after CNO injection, the averaged minute volume for 0-30 min and 30-75 min after morphine injection are used. Values during the episodes when the sensor accidentally fell off due to the animal body rotation were excluded.

#### Stereotaxic surgery

Mice were anesthetized with isoflurane (5% induction, 1.5-2% maintenance with a nose cone; Dräger Vapor 2000, Draegar, Inc., USA) and placed onto a water recirculating heating pad throughout the surgery. Mice were placed on a stereotaxic frame (David Kopf Instruments, USA), the skull was exposed, and the cranium was drilled with a micro motor handpiece drill (Foredom, USA: one or two holes for viral injection(s) or pharmacological delivery, two holes for screws with implantation, one or two holes for optic fibers, and one hole for a micro thermistor). The virus or drug was injected unilaterally (right side) or bilaterally into the PBL (anteroposterior (AP), −1 mm from lambda; mediolateral (ML), ±1.5 mm; dorsoventral (DV), −3.5 mm, re-zero at the midline with the same AP). Injection of the virus or drug was administered with a glass pipette (tips broken for an inner diameter of 20 μm) connected to the Nanoject III Programmable Nanoliter Injector (Drummond Scientific, USA) at a rate of 60 nL/min. Naloxone was injected at a rate of 100 nL/min with a syringe (65458-01, Hamilton, USA) connected to an ultra-micropump (UMP-3, World Precision Instruments, USA). A glass pipette or syringe needle was retracted from the brain slowly after 5 - 10 min. For implantation, optic fibers were implanted above the injection site with the DV noted below, and the micro thermistor head was carefully lowered into the hole above the nasal cavity (AP +3.5 from the nasal fissure, ML 0.3). The implants were covered with superglue and dental cement for stabilization. Behavioral experiments were performed three weeks after virus injection and one week after the micro-thermistor implantation unless otherwise noted.

For the PBL-specific conditional knockout of the *Oprm1* gene, *Oprm1^fl/fl^* mice were bilaterally injected with 200 nL of AAVDJ-CAG-Cre-GFP (1.25E+12 GC/mL) or control AAVDJ-CAG-GFP (2.10E+12 GC/mL) (Salk Institute Viral Vector Core) into the PBL.

For PBL-specific rescue of the *Oprm1* gene, *Oprm1^Cre/Cre^* mice were bilaterally injected with 200 nL of AAV-hSyn-DIO-mCherry-T2A-FLAG-hOprm1 (1.5E+13 GC/mL) or control AAV-DIO-eYFP (2.12E+12 GC/mL) into the PBL.

For fiber photometry, *Oprm1^Cre/+^* mice were unilaterally injected with 200 nL of AAV-DIO-jGCaMP7s (3.75 E+13 GC/mL) or control AAV-DIO-eYFP (2.12E+12 GC/mL) into the PBL, and a stainless steel mono fiberoptic cannula (400 μm diameter, 0.37 NA, Doric Lenses) was implanted 0.05 mm above the injection site.

For chemogenetics, 200 nL of AAV-DIO-hM3Dq-mCherry (6.56E+11 GC/mL), AAV-DIO-KORD-mCitrine, or control AAV-DIO-eYFP (2.12E+12 GC/mL) was bilaterally injected into the PBL of *Oprm1^Cre/+^* (for hM3Dq) or *Oprm1^Cre/Cre^* (for KORD) mice.

#### Pharmacology

Morphine sulfate (Spectrum Chemical, USA) was dissolved in 0.9% saline to make a 4 mg/mL working concentration. The final concentration of morphine was 10 mg/kg in PBL-specific *Oprm1* knockout tests, 80 mg/kg in PBL-specific rescue of OIRD, and 40 mg/kg in all other experiments. For the characterization of morphine-induced respiratory depression, mice were introduced into the WBP chamber 30 - 40 min after morphine injection.

For PBL-specific rescue of OIRD with naloxone, wild-type mice were kept under isoflurane anesthesia until the breathing rate was stable for 10 min. Then, 80 mg/kg morphine (s.c.) was systemically administered. After 30 min, 200 nL of 0.4 mg/mL naloxone (Somerset Therapeutics, USA) and FluoSpheres (540/560, 10% v/v, Thermo Fisher, USA) mixture or a control 0.9% saline-FluoSpheres mixture was stereotaxically injected bilaterally into the PBL. Breathing was analyzed from three 2-min episodes: immediately before morphine injection, 30 min after morphine injection, and 10 min after naloxone or saline injection.

For endogenous receptor activation of PBL^*Oprm1*^ neurons, wild-type mice were kept under 1-1.5% isoflurane anesthesia until breathing rate is stable (fluctuation < 10 bpm) for 10 min. Then, 80 mg/kg morphine (s.c.) was systemically administered. After 30 min, 200 nL of the pharmacological agent (Supplementary Table 1) was stereotaxically injected into the PBL bilaterally. 50 nL of Cholera Toxin Subunit B (CTB)-555 was subsequently injected with the same coordinates to mark the injection site. Breathing was analyzed from three 2-min episodes: immediately before morphine injection, 30 min after morphine injection, and 30 min after drug injection.

#### Fiber photometry

Fiber photometry (1-site Fiber Photometry System, 405 and 465 nm, Doric Lenses Inc, Canada) with Doric Neuroscience Studio software was used to record PBL^*Oprm1*^ neural activities. GCaMP isosbestic fluorescence (405-nm excitation) and calcium-dependent fluorescence (465-nm excitation) were recorded at a sampling rate of 12 kHz, and data were analyzed with the Doric Neuroscience Studio software. F0 was calculated by a least mean squares fitting of the 405-nm channel relative to the 465-nm channel, and ΔF/F was calculated as (F_465_-F_405_fitted_)/F_405_fitted_. Data were further analyzed with custom MATLAB scripts.

For concurrent measurements of PBL^*Oprm1*^ neural activity and respiration, animals were recorded in their home cage for 30 min before and 30 min after intraperitoneal (i.p.) morphine injection at 40 mg/kg. Cross-correlation analysis between the calcium signals and respiration data was performed with the z-scored data, then with the MATLAB “xcorr” function using the normalized option such that autocorrelations at zero lag equal 1. Calcium peaks were detected with the MATLAB “findpeaks” function.

#### Convergent cross-mapping

State-space reconstruction models were generated using the framework of convergent cross mapping (34), a nonlinear time series embedding method (53) based on the Takens theorem and its generalized form (54) that builds low-dimensional manifolds from time series and makes predictions across variables. Analysis and predictions were calculated using the R package rEDM 0.7.2 (https://cran.r-project.org/web/packages/rEDM/) for evaluation and rEDM 0.7.4 (https://ha0ye.github.io/rEDM/) for model predictions in the RStudio environment. This program was run on a dual Intel Xeon Gold 6148 Server with 384GB RAM or an Intel Core i9 2.4 GHz MacBook Pro with 32 GB RAM. Key parameters were determined individually by lagged coordinate embedding using the simplex function implementation in rEDM to optimize predictive skill as measured by the predicted over observed rho. Parameters include the delay tau, which gives the characteristic timescale of the time series, and the embedding dimensionality, which estimates the number of variables driving the system and approximates the real number of variables as given by the Whitney embedding theorem (55) as minimally equal to the real number n of variables, but no more than two times n + 1 (*n* ≤ *E* ≤ 2*n* + 1). The choice of tau was also informed by minimizing mutual information (56). This approximately corresponds to an autocorrelation of ~0.3, which was applied if it maximized predictive skill across datasets. To prevent data contamination, an exclusion radius was applied that was larger than the respiration rate smoothing window of five timesteps. Whenever the data allowed, an exclusion radius of E*tau was applied, unless the data was insufficient to apply this upper bound. In this case, the exclusion radius would be made just larger than tau.

#### Chemogenetics

For KORD-mediated inhibition, salvinorin B (SALB, Cayman Chemical, USA) was diluted in 100% DMSO to make a 15 mg/mL stock solution, then further diluted in 0.9% saline (10% v/v) to make a 1.5 mg/mL working suspension. The control used consisted of DMSO diluted in 0.9% saline to make 10% v/v. Mice were injected with 5 μL/g of body weight. The final concentration was 7.5 mg/kg. Behavioral testing was performed immediately after the SALB injection. Animals that did not show the viral expression on one side were excluded from the analysis.

For hM3Dq-mediated excitation, clozapine-N-oxide (CNO, Cayman Chemical, USA) was diluted in 0.9% saline to make a 1 mg/mL working solution. Mice were injected with 5 μL/g of body weight, and the final concentration was 5 mg/kg. For quantification of the CNO effect, both the 0 - 10 min before and 20 - 30 min after CNO injection periods were analyzed.

#### RiboTag profiling and library generation

Isolation of polysome-associated mRNA using RiboTag was performed as previously described with minor modification (37, 57). Briefly, 250-μm thick slices containing the PBL were obtained using a VT 1200S Vibratome (Leica, Germany). The region of interest was further dissected with surgical scissors. Tissues were transferred into 1.5 mL microcentrifuge tubes containing homogenization buffer (50 mM Tris, pH 7.5, 100 mM KCl, 12 mM MgCl_2_, 1% Nonidet P-40, 1 mM dithiothreitol (DTT), 200 U/mL RNasin, 1 mg/mL heparin, 100 μg/mL cycloheximide, and 1× protease inhibitor mixture) and mechanically dissociated and lysed using pellet pestles (7495211500-DS, DWK Life Sciences LLC, USA). Lysates were centrifugated for 10 min at 12,000 rpm at 4 °C. Post mitochondrial supernatants were transferred to fresh 1.5 mL microcentrifuge tubes. For immunoprecipitation, 4 μL of anti-hemagglutinin1.1 antibody (BioLegend, USA) was added to the lysate-containing tube and incubated for 4 h at 4 °C on a microtube rotator. After incubation, magnetic protein A/G beads (Cat. No. 88803; Thermo Fisher Scientific) were added to the lysate with antibody prior to incubation on a microtube rotator at 4 °C overnight. The following day, the microcentrifuge tubes were placed into the magnetic stand on ice. The supernatant was removed from sample tubes and used for non-Oprm1 control subsequently. The magnetic beads were washed with high-salt buffer (50 mM Tris, pH 7.5, 300 mM KCl, 12 mM MgCl_2_, 1% Nonidet P-40, 1 mM DTT, and 100 μg/mL cycloheximide) to remove the non-specific binding from immunoprecipitation. After washing, 350 μL of RLT plus buffer with β-mercaptoethanol from the RNeasy Mini Kit (Qiagen, Germany) was added. The extraction of total RNA was performed with the RNeasy Mini Kit. All RNA samples were quantified with the *Qubit RNA Assay Kit (Invitrogen, USA) and analyzed* with the RNA 6000 Pico Kit (Agilent, USA).

Isolated RNA was prepared using the Trio RNA-Seq (Cat. No. 0507-08; NuGEN, USA). Briefly, cDNA was synthesized from the total RNA using reverse transcriptase with oligo dT and resynthesized to produce double-stranded cDNA. After amplification of double-stranded cDNA, cDNA was purified with AMPure XP Beads (Cat. No. A63881; Beckman Coulter, USA), fragmented to the library, and classified using a barcoded adaptor. All libraries were quantified by qPCR and analyzed *with the RNA 6000 Pico Kit.*

#### RiboTag profiling analysis

RNA library quality was confirmed using the 2100 Bioanalyzer (Agilent, USA). Barcoded samples *were pooled* and sequenced on the NextSeq500 (Illumina, USA) with the 75-bp read length single-end format. Image analysis and base calling were conducted using the Illumina CASAVA-1.8.2 software. Sequencing read quality was evaluated with the FastQC package (http://www.bioinformatics.babraham.ac.uk/projects/fastqc). Fastq reads were aligned to the reference genome (GRCm38.p6) using the STAR tool (version 2.7.2) (58). The quantification package RSEM (version 1.2.28) (59) was utilized to calculate gene expression from BAM files. In doing so, estimated count and TPM (Transcripts Per Million) were computed. Fold changes were calculated from TPM values (estimated counts > 20) between HA-tag and controls. To visualize fold changes, we utilized the ggplot2 package from R. UP (> 3 fold change) and DOWN (< −3 fold change) were highlighted with orange and blue colors, respectively. Key genes for the parabrachial marker and GPCRs were further annotated.

#### Histology

Mice were euthanized with CO_2_ at a flow rate of 1.2 liters per minute (LPM), perfused intracardially with ice-cold phosphate-buffered saline (PBS), and fixed with 4 % paraformaldehyde (PFA) in phosphate buffer (PB). The brain was extracted, post-fixed in 4 % PFA overnight and dehydrated in 30 % sucrose in PBS until sliced. Frozen brains were cut into 50 μm coronal slices using a CM 1950 cryostat (Leica, Germany) and stored in PBS before mounting. The slices were mounted on Superfrost Microscope Slides (Fisher Scientific, USA) with DAPI Fluoromount-G mounting media (Southern Biotech, USA) for imaging.

#### Immunohistochemistry

To validate the co-expression of Cre:GFP and Rpl22:HA in PBL^*Oprm1*^ neurons, immunohistochemistry for HA and GFP was performed. To validate the conditional knockout of the *Oprm1* gene in the PBL, we performed immunohistochemistry for MOR.

Mice were euthanized with CO_2_ at a flow rate of 1.2 LPM, perfused intracardially with ice-cold PBS, and fixed with 4 % PFA in PB. The brain was extracted, post-fixed in 4 % PFA overnight and dehydrated in 30 % sucrose in PBS until sliced. Frozen brains were cut into 30 μm coronal slices with a CM 1950 cryostat and stored in PBS.

Sections were washed with PBST (PBS with 3% Tween-20, Fisher BioReagents, USA). After blocking with 3 % normal donkey serum (NDS, Jackson ImmunoResearch Inc., USA) for 1 h at room temperature and rinsing with PBST, the slices were incubated with the corresponding antibodies: mouse monoclonal anti-HA1.1 (1:1000, BioLegend, USA), chicken anti-GFP (1:1000, Aves Labs Inc, USA), and rabbit anti-MOR (1:1000, ImmunoStar, USA) antibodies at 4 °C for overnight (anti-HA1.1 and anti-GFP) or 24 h (anti-MOR). The next day, brain tissues were rinsed with PBST and incubated in Alexa Fluor^®^ 647-conjugated Donkey Anti-Mouse IgG, Alexa Fluor^®^ 488-conjugated Donkey Anti-Chicken IgY, and Alexa Fluor^®^ 647-conjugated Donkey Anti-Rabbit IgG (1:1000, Jackson ImmunoResearch Inc., USA) in 3 % NDS for 90 min at room temperature. After washing with PBS, brain slices were mounted on Superfrost Microscope Slides with DAPI Fluoromount-G mounting media for imaging.

#### RNA *in situ* hybridization

RNA *in situ* hybridization was performed using the RNAscope Fluorescent Multiplex Assay using the probes and kits purchased from Advanced Cell Diagnostics (ACD, USA). Brains were collected from wild type mice and immediately frozen with 2-methylbutane chilled with dry ice. Frozen brains were cut into 20-μm coronal slices with a CM 1950 cryostat and directly mounted onto the Superfrost Plus Microscope Slides (Fisher Scientific). Sample preparation, pretreatment, and signal detection were performed according to the ACD protocols. Probes used are listed below: *Oprm1* (ACD #315841), *Htr2a* (#401291), *Cckar* (#313751), *Drd5* (#494411), *Tacr3* (#481671), *Tacr1* (#428781), *Cre* (#402551), *Slc17a6* (#319171), and *Slc32a1* (#319191).

Two to three representative images from the PBL (AP = −5.0 to −5.2) were selected from n = 3 mice, and PBL cells within a field of view of 300 μm x 300 μm (for *Oprm1* colocalization with *Cre*, *Htr2a*, *Cckar*, *Tacr3*, *Tacr1*, and *Drd5*) or 600 μm x 600 μm (for *Oprm1* colocalization with *Slc17a6* and *Slc32a1*) were used for quantification. Quantification of the colocalization is done manually with ImageJ software according to the ACDBio technical note. DAPI-stained nuclei were first identified, then the cell contour was defined with a 2-μm radius surrounding the DAPI signals. Cells containing at least five puncta inside the imaginary boundary were labeled as positive.

#### Imaging

Images for verification of injection and implantation site and anti-MOR staining for conditional knockout validation were taken with a BZ-X710 all-in-one fluorescence microscope with BZ-X viewer software under a 10X, 0.45 NA objective (Keyence, Japan). Images for anti-HA & anti-GFP immunostaining, RNAscope, and *Oprm1^Cre^*::Ai14 expression characterization were acquired with an FV3000 Confocal Laser Scanning Microscope with FV31S-SW software under a 20X, 0.75 NA or 40X, 0.95 NA UPLSAPO objective (Olympus, Japan). For comparison, images were processed with the same gain, offset, and exposure time.

#### Statistical Analysis

All data are shown as mean ± SEM and analyzed using either a student’s t-test, one-way ANOVA with Tukey’s *post hoc* comparison, or two-way ANOVA with Bonferroni’s *post hoc* comparison. All statistical analyses were performed with Prism 6 (GraphPad Software Inc., USA). ns *p* > 0.05, * *p* < 0.05, ** *p* < 0.01, *** *p* < 0.001, **** *p* < 0.0001.

For a detailed list of resources, please refer to Supplementary Table 2.

## Supplementary Figures

**Supplementary Figure 1.**
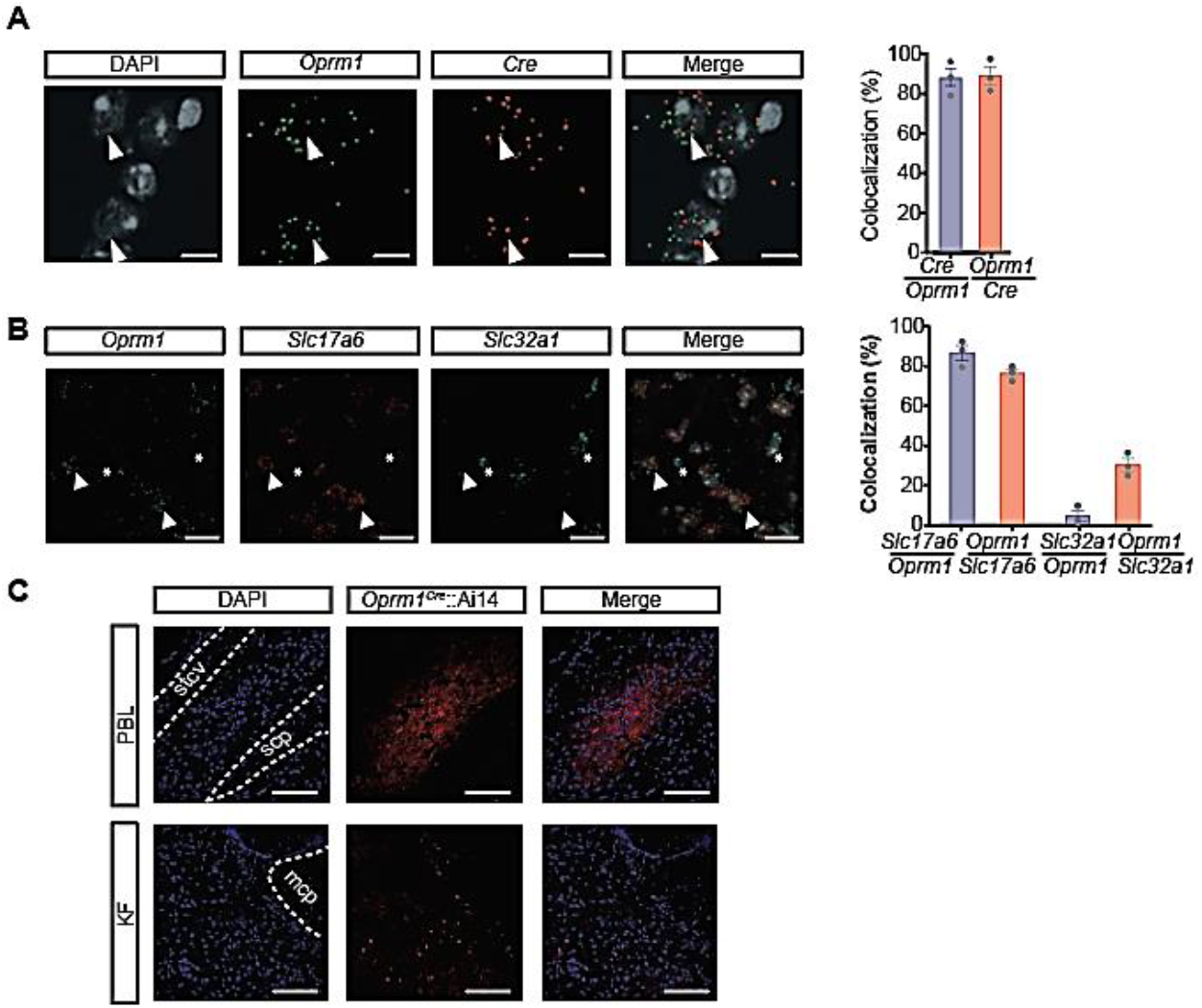
Characterization of the *Oprm1^Cre^* mouse line and PBL^*Oprm1*^ neurons. (**A**) RNA *in situ* hybridization and quantification of the colocalization for *Oprm1* and *Cre* mRNAs in the PBL of *Oprm1^Cre/+^* mice (n = 902 cells from 3 animals). Arrowheads, double-labeled cells. Scale bar, 10 μm. (**B**) RNA *in situ* hybridization and quantification of the colocalization for *Oprm1*, *Slc17a6* (encoding VGLUT2), and *Slc32a1* (encoding VGAT) mRNAs in the PBL of wild type mice (n = 1552 and 1431 cells from 3 animals). Arrowheads, *Oprm1^+^ / Slc17a6^+^* cells; asterisks, *Oprm1^−^ / Slc32a1^+^* cells. Scale bar, 50 μm. (**C**) Example histology from *Oprm1^Cre^::*Ai14 double transgenic mice showing tdTomato expression in the PBL and KF under the same microscope settings. Anatomical landmarks for PBL: stcv, ventral spinocerebellar tract; scp, superior cerebellar peduncle; mcp, middle cerebellar peduncle. Scale bar, 200 μm.

**Supplementary Figure 2.**
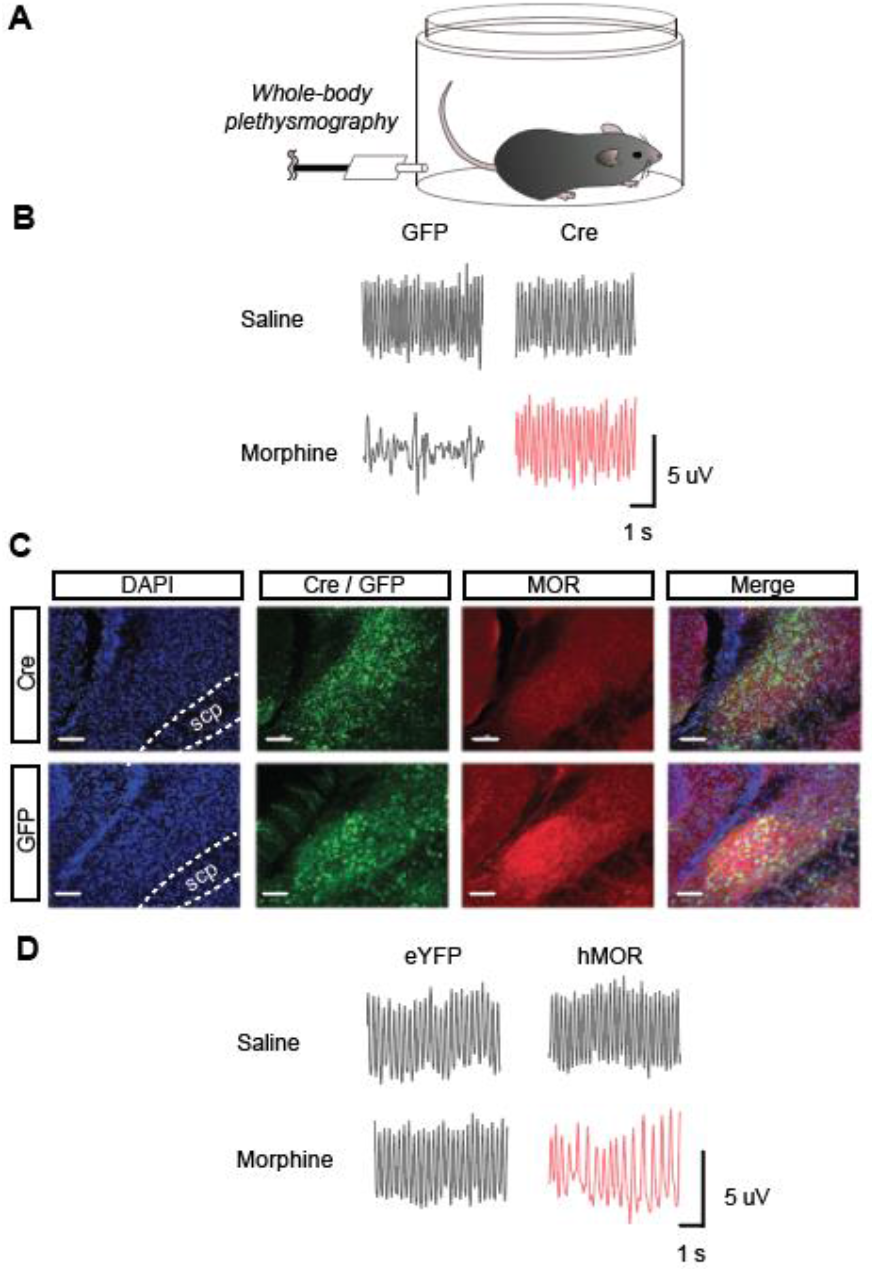
MOR signaling in the PBL is indispensable for OIRD. (**A**) Whole-body plethysmography measured respiratory parameters before and after systemic injection of saline and morphine, after conditional ablation (**B**) and rescue (**D**) of the MOR signaling in the PBL. (**B**) Example plethysmograph after saline and morphine injections into the *Oprm1^fl/fl^* mice expressing AAV-GFP and AAV-Cre-GFP in the PBL. (**C**) Confirmation of MOR deletion by MOR immunohistochemistry after stereotaxic injection of AAV-Cre-GFP and control AAV-GFP into the PBL of the *Oprm1^fl/fl^* mice. Scale bar, 100 μm. (**D**) Example plethysmograph after saline and morphine injections into the *Oprm1^Cre/Cre^* mice expressing AAV-DIO-eYFP and AAV-DIO-hMOR in the PBL.

**Supplementary Figure 3.**
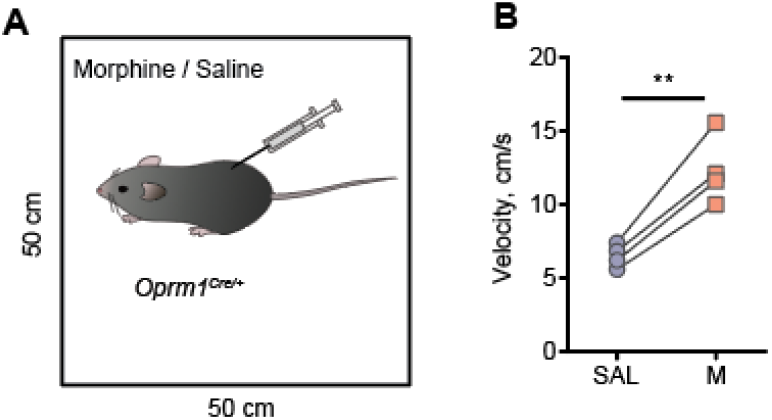
Morphine effect on locomotor sensitization. (**A**) Locomotor activity of *Oprm1^Cre/+^* mice after morphine (40 mg/kg, i.p.) or saline injection was measured in the open field. (**B**) Systemic morphine injection increased locomotor activity in *Oprm1^Cre/+^* mice. Paired t-test, **, *p* < 0.01.

**Supplementary Figure 4.**
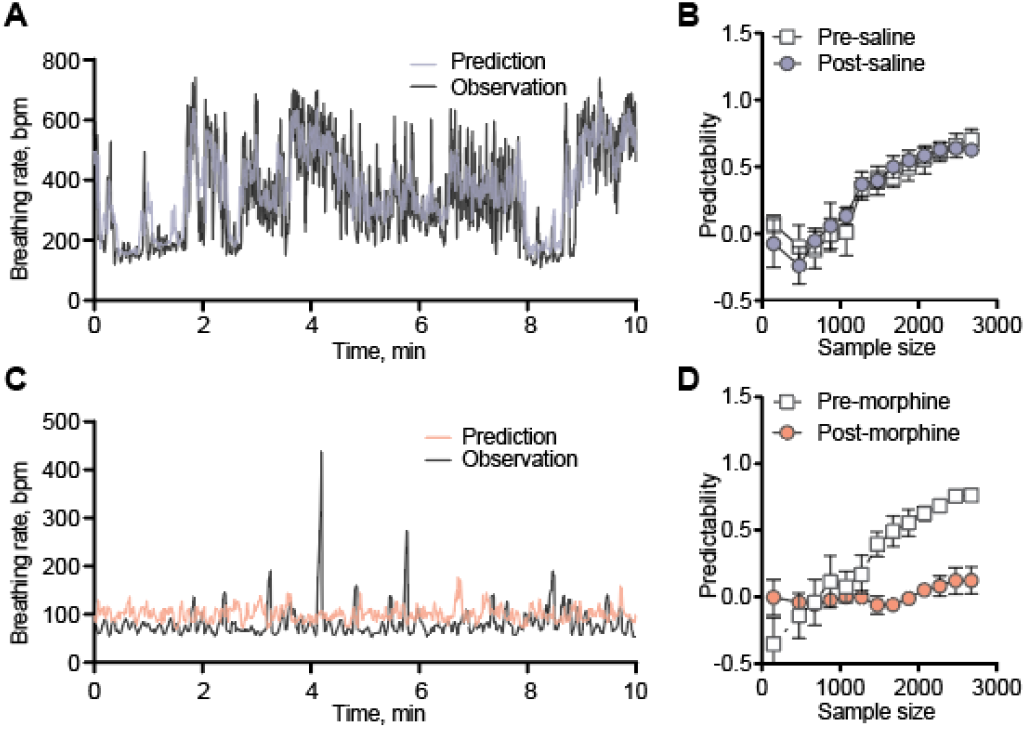
Morphine eliminated the breathing predictor. Convergent cross-mapping (CCM) prediction of breathing rate using calcium activity as inputs before and after saline (0.9%, i.p., **A** and **B**) and morphine (40 mg/kg, i.p., **C** and **D**) injection. (**A**) After saline injection, the predicted and observed breathing rate traces followed closely with each other during the 10-min example. (**B**) Model predictability increased with the sample size before and after saline injection (n = 4). (**C**) After morphine injection, the predicted and observed breathing rate traces no longer matched with each other. (**D**) Model predictability increased with the sample size before morphine injection, but the relationship was completely abolished after morphine injection (n = 4).

**Supplementary Figure 5.**
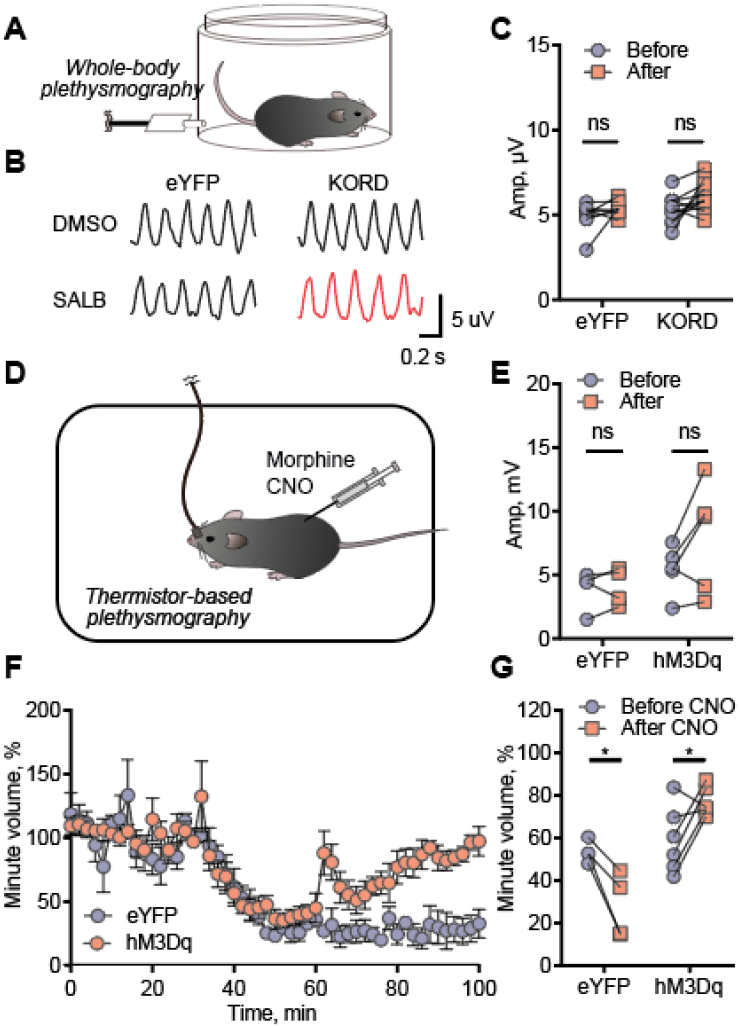
(**A**) Whole-body plethysmography was used to measure respiratory parameters before and after systemic injection of DMSO and SALB in the *Oprm1^Cre/Cre^* mice expressing AAV-DIO-eYFP and AAV-DIO-KORD in PBL^*Oprm1*^ neurons. (**B**) Example plethysmograph after DMSO and 7.5 mg/kg SALB injections in eYFP and KORD groups. KORD-expressing animals displayed a slower respiratory rate after SALB injection compared to other groups. (**C**) Respiratory amplitude was not significantly changed before and after SALB injection, in both eYFP (n = 8) and KORD (n = 11)-expressing animals. Two-way ANOVA with Bonferroni’s multiple comparison post-hoc test, ns, not significant. (**D**) Thermistor-based plethysmography was used for measuring respiration before and after systemic injection of 40 mg/kg morphine and CNO in the *Oprm1^Cre/+^* mice expressing AAV-DIO-eYFP and AAV-DIO-hM3Dq in PBL^*Oprm1*^ neurons. (**E**) Respiratory amplitude was not significantly changed before and after CNO injection, in both eYFP (n = 5) and hM3Dq-expressing animals (n = 6). Although there was a trend of increase in the hM3Dq group, it failed to reach statistical significance. Two-way ANOVA with Bonferroni’s multiple comparison post-hoc tests, ns, not significant. (**F**) Activation of PBL^*Oprm1*^ neurons by injecting CNO (5 mg/kg, i.p.) in the hM3Dq-expressing group (n = 6) completely rescued the minute volume to baseline level after morphine-induced respiratory depression, but not in eYFP-expressing group (n = 5). (**G**) Quantitative analysis of **F** showing CNO injection significantly increased the minute volume in the hM3Dq-expressing group (n = 6), whereas failed to rescue in the eYFP-expressing group (n = 5). Two-way ANOVA with Bonferroni’s multiple comparison post-hoc test. *, *p* < 0.05.

**Supplementary Figure 6.**
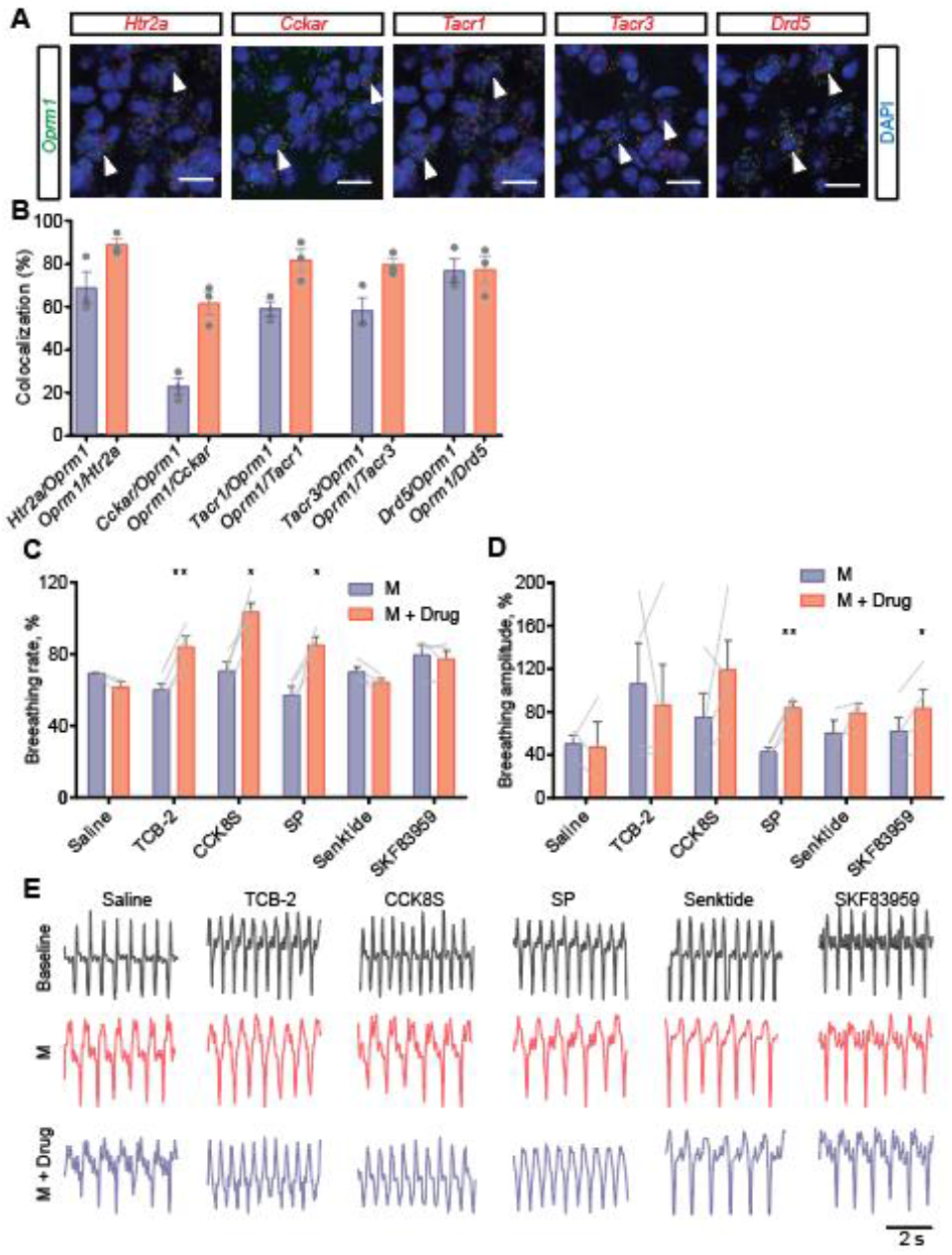
Rescuing of OIRD by activating endogenous GPCRs expressed on PBL^*Oprm1*^ neurons. (**A**) RNA *in situ* hybridization confirms the co-expression of mRNA of *Oprm1* and five selected GPCRs, *Htr2a*, *Cckar*, *Tacr1*, *Tacr3*, and *Drd5*, in the wild type mice. Arrowheads, double-labeled cells. Scale bar, 50 μm. (**B**) Quantification of RNA *in situ* hybridization showing the colocalization of *Oprm1* and selective GPCR genes in the PBL of wild type mice (770, 539, 836, 657, 431 cells for *Htr2a*, *Cckar*, *Tacr1*, *Tacr3*, and *Drd5*, respectively). (**C-D**) Normalized breathing rate (**C**) and amplitude (**D**) before morphine injection (Baseline), 30 minutes after morphine injection (80 mg/kg, s.c., M), and 30 minutes after drug injection into the PBL (M + Drug), for all six drugs tested. Large variations in amplitude are due to the technical limitations of the piezoelectric sensor since its location is subjective to the slight movements of the animal’s body. Paired t-test, *, *p* < 0.05; **, *p* < 0.01. **e** Example plethysmograph showing the breathing rhythm changes at each stage of the experiment. Note the decreased breathing rate after morphine injection and increased breathing rate after TCB-2, CCK8S, and SP injections. The exact shape of the plethysmograph varies due to the placement of the piezoelectric sensor.

**Supplementary Figure 7.**
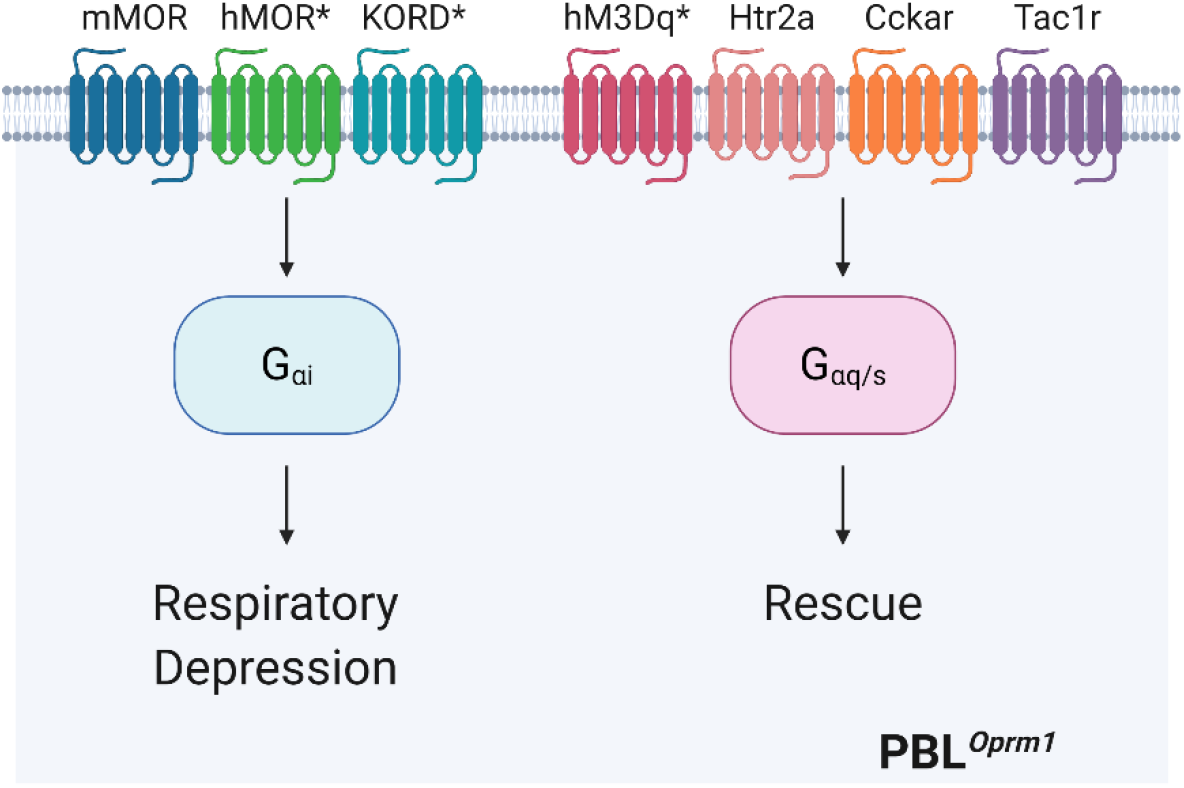
Summary of the current study. PBL^*Oprm1*^ neurons are critical players in OIRD pathogenesis and promising therapeutic targets for treating OIRD. In intact mice, inhibition of PBL^*Oprm1*^ neurons through G_i_-coupled GPCRs via endogenous MOR (mMOR), human MOR (hMOR), and KOR-derived DREADD (KORD) leads to respiratory depression. In contrast, activation of these neurons via artificial (hM3Dq) or endogenous G_q/s_-coupled GPCRs (Htr2a, Cckar, Tac1r) rescues OIRD. Artificial GPCRs are marked with asterisks. Created with BioRender.com.

**Supplementary Table 1.** List of pharmacological agents used in the current study.

| Drug Name | Receptor (Gene) | Receptor Family | Concentration | Solvent | Company |
| --- | --- | --- | --- | --- | --- |
| Naloxone | MOR (*Oprm1*) | Gi | 0.4 mg/mL (i.c., intracranial) | 0.9% saline | Somerset Therapeutics |
| Morphine | MOR (*Oprm1*) | Gi | 80 mg/kg (s.c., for anesthesia experiments);<br>10 mg/kg (i.p., for PBL-specific *Oprm1* knockout);<br>40 mg/kg (i.p., for all other experiments) | 0.9% saline | Spectrum Chemical |
| Salvinorin B (SALB) | KORD | Gi | 7.5 mg/kg (i.p.) | 0.9% saline containing 10% DMSO | Cayman Chemical |
| Clozapine-N-oxide (CNO) | hM3Dq | Gq | 5 mg/kg (i.p.) | 0.9% saline | Cayman Chemical |
| TCB-2 | 5-HT<sub>2A</sub> (*Htr2a*) | Gq | 1 mg/mL (i.c.) | 0.9% saline | Tocris Bioscience |
| CCK Octapeptide, sulfated (CCK8S) | CCK<sub>1</sub>R (*Cckar*) | Gq | 1 mg/mL (i.c.) | 0.9% saline | Abcam |
| Substance P | NK<sub>1</sub>R (*Tacr1*) | Gq, Gs | 1 mg/mL (i.c.) | 0.9% saline | Cayman Chemical |
| Senktide | NK<sub>3</sub>R (*Tacr3*) | Gq | 1 mg/mL (i.c.) | 0.9% saline | Cayman Chemical |
| SKF-83959 | D<sub>5</sub>R (*Drd5*) | Gs | 1 mg/mL (i.c.) | 0.9% saline containing 5% DMSO | Tocris Bioscience |

**Supplementary Table 2.** Key Resources Table.

| Type | Designation | Source or reference | Identifiers |
| --- | --- | --- | --- |
| Mouse strains | Oprm1-Cre:GFP | Laboratory of Dr. Richard Palmiter | N/A |
|  | Wild type C57BL/6J | Jackson Laboratory | Stock No.<br>000664 |
|  | RiboTag *Rpl22<sup>HA/HA</sup>* | Jackson Laboratory | Stock No.<br>011029 |
|  | Ai14 *Gt(ROSA)26Sor<sup>tm14(CAG-tdTomato)Hze</sup>* | Jackson Laboratory | Stock No.<br>007914 |
| Virus | AAVDJ-CAG-Cre-GFP | Salk Institute Viral Vector Core | N/A |
|  | AAVDJ-CAG-GFP | Salk Institute Viral Vector Core | N/A |
|  | AAV-hSyn-DIO-mCherry-T2A-FLAG-hOprm1 | Laboratory of Dr. Matthew Banghart | N/A |
|  | AAV1-syn-FLEX-jGCaMP7s-WPRE | Addgene | Addgene#<br>104491 |
|  | AAVDJ-EF1a-DIO-hM3D(Gq)-mCherry | Salk Institute Viral Vector Core | Addgene#<br>50460 |
|  | AAV-DIO-KORD-mCitrine | Laboratory of Dr. Richard Palmiter | N/A |
|  | AAV1-DIO-eYFP | Laboratory of Dr. Richard Palmiter | N/A |
| Antibodies | Anti-hemagglutinin1.1, mouse (1:1000) | BioLegend | RRID<br>AB_2565335 |
|  | Anti-GFP, chicken (1:1000) | Aves Labs | RRID<br>AB_2307313 |
|  | Anti-MOR, rabbit (1:1000) | ImmunoStar | RRID<br>AB_572251 |
|  | Alexa Fluor® 647-conjugated Donkey Anti-Mouse IgG (1:1000) | Jackson ImmunoResearch Laboratories | RRID<br>AB_2340863 |
|  | Alexa Fluor® 488-conjugated Donkey Anti-Chicken IgY (1:1000) | Jackson ImmunoResearch Laboratories | RRID<br>AB_2340375 |
|  | Alexa Fluor® 647-conjugated Donkey Anti-Rabbit IgG (1:1000) | Jackson ImmunoResearch Laboratories | RRID<br>AB_2492288 |
| RNAscope probes | Oprm1 | Advanced Cell Diagnostics | 315841 |
|  | Htr2a | Advanced Cell Diagnostics | 401291 |
|  | Cckar | Advanced Cell Diagnostics | 313751 |
|  | Drd5 | Advanced Cell Diagnostics | 494411 |
|  | Tacr3 | Advanced Cell Diagnostics | 481671 |
|  | Tacr1 | Advanced Cell Diagnostics | 428781 |
|  | Cre | Advanced Cell Diagnostics | 402551 |
|  | Slc17a6 | Advanced Cell Diagnostics | 319171 |
|  | Slc32a1 | Advanced Cell Diagnostics | 319191 |
| Chemicals | FluoSpheres 540/560 (10% v/v) | Thermo Fisher | Cat# F8809 |
|  | Cholera Toxin Subunit B-555 | Invitrogen | Cat# C34776 |
|  | Naloxone | Somerset Therapeutics | Cat#<br>70069007110 |
|  | Morphine | Spectrum Chemical | Cat# M1167 |
|  | TCB-2 | Tocris Bioscience | Cat# 2592 |
|  | CCK Octapeptide, sulfated | Abcam | Cat# ab120209 |
|  | Substance P | Cayman Chemical | Cat# 24035 |
|  | Senktide | Cayman Chemical | Cat# 16721 |
|  | SKF-83959 | Tocris Bioscience | Cat# 2074 |
|  | Clozapine N-oxide | Cayman Chemical | Cat# 16882 |
|  | Salvinorin B | Cayman Chemical | Cat# 11488 |
|  | 2-methylbutane | Fisher Chemical | Cat# O35514 |
| Software | Doric Neuroscience Studio | Doric Lenses | N/A |
|  | RStudio | RStudio | Version<br>1.2.5001 |
|  | rEDM | https://cran.r-project.org/web/packages/rEDM/ | Version 0.7.2 |
|  | rEDM | https://ha0ye.github.io/rEDM/ | Version 0.7.4 |
|  | LabChart | ADInstruments | Version 8 |
|  | BZ-X viewer | Keyence | N/A |
|  | OlyVIA | Olympus Life Science | N/A |
|  | PRISM | GraphPad Software | Version 6 |
|  | Illustrator | Adobe | Version CS6<br>and CC2018 |

